# An essential post-developmental role for Lis1 in mice

**DOI:** 10.1101/212340

**Authors:** Timothy J. Hines, Xu Gao, Subhshri Sahu, Meghann M. Lange, Jill R. Turner, Jeffery L. Twiss, Deanna S. Smith

## Abstract

*LIS1* mutations cause lissencephaly (LIS), a severe developmental brain malformation. Much less is known about its role in the mature nervous system. LIS1 regulates the microtubule motor cytoplasmic dynein 1 (dynein), and as LIS1 and dynein are both expressed in the adult nervous system, Lis1 could potentially regulate dynein-dependent processes such as axonal transport. We therefore knocked out Lis1 in adult mice using tamoxifen-induced, Cre-ER-mediated recombination. When an actin promoter was used to drive Cre-ER expression (Act-Cre-ER), heterozygous Lis1 KO caused no obvious change in viability or behavior, despite evidence of widespread recombination by a Cre reporter three weeks after tamoxifen exposure. In contrast, homozygous Lis1 KO caused the rapid onset of neurological symptoms in both male and female mice. One tamoxifen-dosing regimen caused prominent recombination in the midbrain/hindbrain, PNS, and cardiac/skeletal muscle within a week; these mice developed severe symptoms in that time frame and were euthanized. A different tamoxifen regimen resulted in delayed recombination in midbrain/hindbrain, but not in other tissues, and also delayed the onset of symptoms. This indicates that Lis1 loss in the midbrain/hindbrain causes the severe phenotype. In support of this, brainstem regions known to house cardiorespiratory centers showed signs of axonal dysfunction in KO animals. Transport defects, neurofilament alterations, and varicosities were observed in axons in cultured DRG neurons from KO animals. Because no symptoms were observed when a cardiac specific Cre-ER promoter was used, we propose a vital role for Lis1 in autonomic neurons and implicate defective axonal transport in the KO phenotype.

**SIGNIFICANCE STATEMENT:** Mammalian Lis1 is best known for its role in brain development. Lis1 binds to and regulates the microtubule motor, cytoplasmic dynein. We show that Lis1 function is needed post-developmentally and provide evidence that loss of Lis1 in the hindbrain leads to death. The effect is dose dependent in mice, as loss of only one allele does not produce an overt phenotype. However, since LIS1 haploinsufficiency causes lissencephaly (LIS) in humans, our study raises the possibility that post-developmental axonal transport defects could contribute to worsening symptoms in children with LIS1 mutations. Our data are consistent with the hypothesis is that Lis1 regulates dynein-dependent axon transport in the mature nervous system.

## INTRODUCTION

*LIS1* mutations in humans cause a “smooth brain” malformation called lissencephaly (LIS) characterized by severe cognitive and motor impairments and worsening epilepsy, leading to early mortality (Dobyns et al., 1993; Sapir et al., 1999; Gleeson, 2000; Sicca et al., 2003; Saillour et al., 2009; Reiner and Sapir, 2013; Dobyns and Das, 2014; Herbst et al., 2016). Most of the human mutations result in a null allele with ~50% reduction of LIS1 protein levels, which profoundly impacts the developing nervous system. Other mutations can produce a milder phenotype, but the phenotype/genotype correlation is complex. A classic mouse study made it clear that gene dosage is relevant, as progressive reduction of Lis1 protein levels caused progressively more severe phenotypes (Hirotsune et al., 1998). Deletion of a large portion of one Lis1 allele in mice, resulting in a null allele, delays neuronal migration and differentiation, but unlike humans, mature mice show mild neurological abnormalities and are viable and fertile (Hirotsune et al., 1998; Gambello et al., 2003). Cre-mediated knockout (KO) in specific subpopulations of developing neural cells in mice impacts mitosis and nucleokinesis, causing developmental delay (Tsai et al., 2005; Tsai et al., 2007; Yingling et al., 2008; Youn et al., 2009; Hippenmeyer et al., 2010).

Lis1 is a highly conserved regulator of the minus-end directed microtubule motor protein, cytoplasmic dynein 1; together they regulate neural stem cell spindle orientation, nucleokinesis, and nuclear envelope breakdown during brain development (Vallee et al., 2001; Wynshaw-Boris and Gambello, 2001; Gambello et al., 2003; Shu et al., 2004; Tsai et al., 2005; Vallee and Tsai, 2006; Tsai et al., 2007; Hebbar et al., 2008; Schwamborn and Knoblich, 2008; Yingling et al., 2008; Youn et al., 2009; Hippenmeyer et al., 2010; Moon et al., 2014). In fact, mutations in the dynein heavy chain gene *DYNC1H1* can also cause cortical malformations in humans (Vissers et al., 2010; Willemsen et al., 2012; Poirier et al., 2013).

Of particular interest are reports that DYNC1H1 mutations cause later onset neurological disorders, including forms of spinal muscular atrophy (SMA) and Charcot-Marie-Tooth Disease (Weedon et al., 2011; Harms et al., 2012). Additionally, mutations in genes encoding two other dynein regulators DCTN1 and BICD2, cause Perry Syndrome and SMA (Rees et al., 1976; Wider and Wszolek, 2008; Neveling et al., 2013; Oates et al., 2013; Peeters et al., 2013). The extent to which Lis1 functions post-developmentally, especially in minimally proliferative tissues like adult brain, has not been studied extensively. Baraban et al (2012) found that heterozygous *Lis1* KO in 6-week old mice altered synaptic function in the hippocampus in the absence of altered laminar granule cell architecture, but the mechanisms underlying the altered activity are not known (Hunt et al., 2012). It has been shown that Lis1 manipulations impact dynein-dependent axon transport in sensory neurons cultured from adult rats (Smith et al., 2000; Pandey and Smith, 2011). These results suggest that Lis1 is a positive regulator of dynein-based axon transport in adult mammals. This was also found in adult mouse DRG neurons (Klinman and Holzbaur, 2015). Although axon transport studies suggest a role for Lis1 in cultured adult neurons, these neurons do not form synaptic connections, so its involvement in synapse formation and maturation is currently unknown. If Lis1 indeed regulates axon transport in the mature nervous system, Lis1 mutations could have deleterious effects on circuitry in mature brains. We have addressed this fundamental question using a tamoxifen-inducible Cre-Lox system to disrupt Lis1 selectively in adult mice. We show that Lis1 is indispensable in adult mice, and describe unexpected temporal and spatial recombination patterns and how they impact the phenotype of Lis1 KO in adult animals. Our data point to a vital role for Lis1 in cardiorespiratory nuclei in the hindbrain.

## MATERIALS AND METHODS

### Mice

All animal experiments were conducted under a protocol approved by the Animal Care and Use Committee of the [Author University]. Both males and females were used in experiments, and no differences were observed in outcomes between males and females. Four mouse strains (Jackson Laboratories) were used to generate inducible Lis1 KO mice (Table 1, below). **1)** *129S-Pafah1b1^tm2Awb^/J* (Jackson Laboratory 008002, RRID:IMSR_JAX:008002): *loxP* sites flank exons 3-6. Homozygous mice are viable and fertile, but have mild hippocampal abnormalities and express ~75% of WT Lis1 levels (Hirotsune et al., 1998); **2)** *Tg(CAG-cre/Esr1*)5Amc/J* (Jackson Laboratory 004453, RRID:IMSR_JAX:004453) – a chicken β actin promotor drives expression of Cre recombinase fused to a modified estrogen receptor; **3)** *Tg(Myh6-cre/Esr1*)1Jmk/J* (The Jackson Laboratory 005650, RRID:IMSR_JAX:005650) – expression of Cre-ER is under the control of a cardiac-specific α- myosin heavy chain promoter so that tamoxifen stimulates recombination only in cardiac cells; and **4)** *Gt(ROSA)26Sor^tm4(ACTB-tdTomato,-EGFP)Luo^/J* (The Jackson Laboratory 007576, RRID:IMSR_JAX:007576) – a Cre reporter mouse with *loxP* sites flanking a membrane-targeted tdTomato cassette that is positioned upstream of a membrane-targeted EGFP cassette. In the reporter mice, cells exhibit membrane-associated red fluorescence until the tdTomato cassette is deleted by Cre recombinase for expression of membrane-associated EGFP fluorescence, thereby allowing visualization of both recombined and non-recombined cells (Muzumdar et al., 2007).

**Table 1:** Mouse strains and crosses used in these studies

| Founder mouse lines |  |
| --- | --- |
| Jackson Labs strains | Descriptive name used in paper |
| 129S-Pafah1b1<sup>tm2Awb</sup>/J | Lis1<sup>LoxP/+</sup> or Lis1<sup>LoxP/LoxP</sup> |
| Tg(CAG-cre/Esr1*)1Jmk/J | Act-Cre-ER (heterozygous) |
| Tg(Myh6-cre/Esr1*) 1Jmk/J | Myh6-Cre-ER (heterozygous) |
| Gt(ROSA)26Sor<sup>tm4</sup>(ACTB-tdTomato, -EGFP)<sup>Luo</sup>/J | Cre Reporter (heterozygous) |
| All strains below are homozygous for the Cre Reporter |  |
| Lis1 KO strains | Descriptive name used in paper |
| Lis1<sup>LoxP/LoxP</sup> x Act-Cre-ER (het) | Lis1 KO |
| Lis1<sup>LoxP/LoxP</sup> x Myh6-Cre-ER (het) | Myh6 KO |
| Control strains | Descriptive name used in paper |
| Act-Cre-ER | No flLis1 Control |
| Lis1<sup>LoxP/LoxP</sup> | No Cre Control |
| Lis1<sup>LoxP/+</sup> x Act-Cre-ER | Lis1 KO Het |

Table 1 shows the crosses that were used to generate experimental animals, and shows the descriptive names used for each throughout the manuscript. All strains used in experiments were homozygous for the Cre reporter. Genotyping of all animals was performed using primers and protocols recommended by Jackson Laboratories. Primers are available on request.

### Tamoxifen administration

Tamoxifen was delivered by intraperitoneal (IP) or intracerebroventricular (ICV) injections in adult mice (2-5 months old). For IP delivery, mice were injected with 40 mg/ml tamoxifen (Sigma-Aldrich) dissolved in 10% ethanol and 90% corn oil (Sigma). Two different daily tamoxifen dosage regimens were tested based a previous study using this Lis1 KO strain (Hayashi and McMahon, 2002): Regimen 1 (5 × 2 mg) - 2 mg for five consecutive days (total 10 mg); and, Regimen 2 (2 × 8 mg) - 8 mg injected on two consecutive days (total 16 mg). For ICV delivery, mice were anesthetized with an isoflurane/oxygen vapor mixture (5% induction, 2-3% maintenance) and placed in a stereotaxic device (Kopf Instruments, Tujunga, CA). 5 µl of 50 mM (Z) 4-hydroxytamoxifen (Sigma) dissolved in 100% ethanol was infused into the left lateral ventricle at a rate of 0.4 µl/min using a 5 µl Hamilton syringe. The needle was left in place for one additional min before removal to allow for diffusion from the injection site. Coordinates (-1.0 mm posterior from Bregma, ± 1.0 mm mediolateral, and -2.5mm ventral to skull surface) were determined using the atlas of Paxinos and Franklin (Franklin and Paxinos).

### Analysis of Cre-mediated recombination by tdTomato/EGFP fluorescence

Animals under deep isofluorane anesthesia were perfused transcardially with ice-cold PBS, followed by 4% paraformaldehyde (PFA) in 0.1 M PBS [pH 7.4]. Prior to sectioning, tissues were cryoprotected by immersion overnight in 15% sucrose, followed by 24 hours in 30% sucrose in PBS. Tissues were then frozen in OCT compound (Fisher) using a beaker of 2-methylbutane chilled in dry ice. 10 or 50 µm thick cryosections were stored at -80°C until use. Whole brains and hearts were imaged immediately after dissection using an Olympus SZX-12 with an SZX-RFL2 coaxial fluorescence attachment. Cryosections were imaged using a Leica TCS SP8X confocal microscope equipped with LAS X software and a 63X oil immersion objective (1.4 N/A). Some images were obtained using a Zeiss Axiovert 200 inverted microscope equipped with AxioVision software and a Plan-Neo 100Å~/1.30 and Plan-Apo 63Å~/1.40 oil-immersion objectives (Immersol 518F; Carl Zeiss, Inc.) or a Plan-Neofluor 20x dry objective.

### Protein isolation and immunoblotting

Tissues were dissected quickly from CO_2_-euthanized mice and frozen in liquid nitrogen, followed by Dounce homogenization in ice-cold RIPA lysis buffer with protease and phosphatase inhibitors (Thermo). Total protein in extracts was determined using a BCA assay (Thermo). Automated capillary electrophoresis and immunoblotting (Figures 1 and 2) was performed with the Wes Simple Western system using the manufacturer’s protocol (Protein Simple). For this, 1 µg of lysate was loaded for each sample. Anti-mouse (ERK1), anti-rabbit (Lis1), and total protein detection modules were used per manufacturer’s instructions. Blots were analyzed using Compass Software (Protein Simple). For traditional western blotting (Figures 4 and 7) 10 µg of each sample were separated on 10% acrylamide gels, then transferred to PVDF membrane. Blots were probed with antibodies against Lis1 and dynein intermediate chain and proteins detected by chemiluminescence.

**Figure 1.**
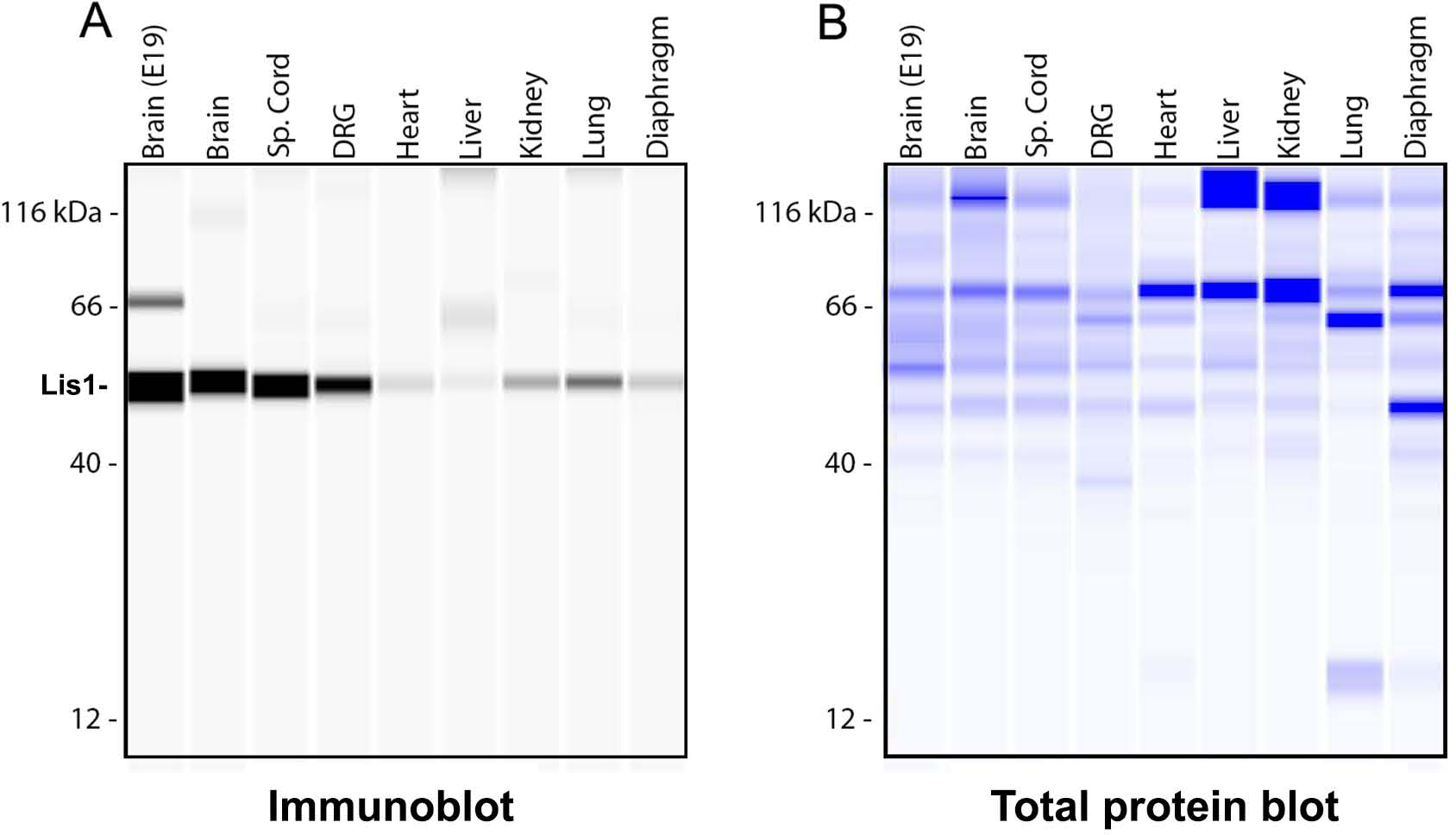
Lis1 protein is expressed in adult mouse tissues. 1 µg of tissue lysates were analyzed using the Wes Simple Western System. Brain extracts from E19 were loaded as a positive control. All other extracts are from 2-month old animals. The size-based separation is processed by Compass software and displayed as virtual blots/gels. **A)** Immune detection of Lis1 in protein samples, depicted in a virtual immunoblot generated by the system. **B)** Total protein detection, visualized by a virtual Coomassie gel generated by the system.

**Figure 2.**
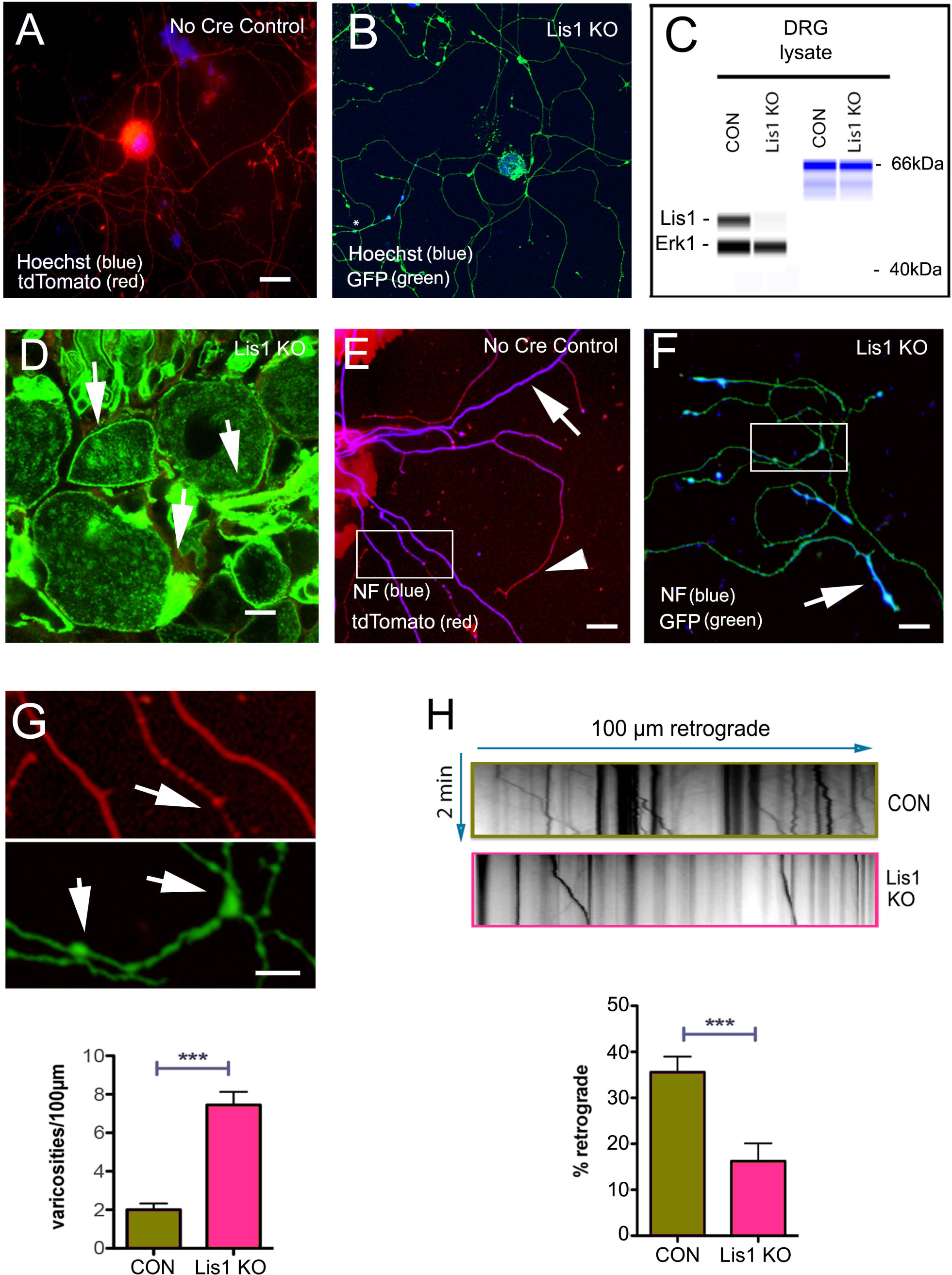
Lis1 KO impacts axonal function in adult mouse DRG neurons. **A)** Cultured DRG neurons from no cre control exposed to 4-OH tamoxifen for 5 days expressed only tdTomato (red) showing no signs of recombination. **B)** In contrast, Lis1 KO neurons showed strong GFP expression (green) demonstrating recombination. **C)** 4-OH tamoxifen also reduced Lis1 protein levels in Lis1 KO neurons relative to no Cre control neurons (CON). **D)** IP injection of 2 × 8 mg tamoxifen in Lis1 KO mice resulted in GFP expression in intact DRGs after 4 days. Arrows point to DRG plasma membranes. **E)** Cultured DRG neurons prepared from IP injected, no Cre control animals expressed only tdTomato (red). NF (blue) was prominent along axon shafts (white arrow) with less prominent in axon terminals (arrowhead). **F)** DRG neurons prepared from IP injected Lis1 KO mice continued to express GFP (green) in culture, and NF (blue) was most prominent in distal axons and enriched in in varicosities (arrow, F). **G)** Insets from E and F have been digitally enlarged to show axonal varicosities (arrows). The bar graph in G shows the average number of varicosities per 100 µm of axon for Lis1 KO and no Cre control (CON) cultures. **H)** Kymographs were generated from time-lapse movies of LysoTracker labeled organelles in GFP-positive axons. The bar graph shows the percentage moving retrogradely in Lis1 KO and no flLis1 control cultures (CON). Bars in G and H indicate mean +/- 95% CI. Significance determined by Mann-Whitney test (G), students t-test (H), *** p=<0.001. For details see Results. Scale bars in A, D, E – 20 µm; Scale bar in B – 5 µm.

### Sciatic nerve transmission electron microscopy

While anesthetized with isoflurane, WT mice were perfused with PBS, then buffered 2.5% glutaraldehyde. Nerves were removed and fixed overnight in 2.5% glutaraldehyde, then section and stained with osmium tetroxide for imaging on a JEOL 200CX Transmission Electron Microscope.

### Antibodies

Primary antibodies used are as follows: Lis1 rabbit polyclonal 484/485 (Smith et al., 2000) (WB: 1:500), Lis1 rabbit polyclonal (Wes: 1:25, WB: 1:500; Santa Cruz Biotechnology sc-15319, RRID:AB_2159891); ERK1 rabbit polyclonal (Wes: 1:100; Abcam ab109282, RRID:AB_10862274); a mix of the pan-axonal neurofilament mouse monoclonal cocktail (IF: 1:500; BioLegend 837904, RRID:AB_2566782) and the neurofilament 200 kDa mouse monoclonal, clone RT97 (IF: 1:500; Millipore CBL212, RRID:AB_93408) were used to label neurofilaments; choline acetyltransferase goat polyclonal (IF: 1:100; Millipore AB144P, RRID:AB_11214092); MAP2 chicken polyclonal (IF: 1:100; Abcam ab5392, RRID:AB_2138153); myelin basic protein chicken polyclonal (IF: 1:500; Aves MBP, RRID:AB_2313550); α-tubulin mouse monoclonal (WB: 1:2000; Sigma-Aldrich T5168, RRID:AB_477579); dynein intermediate chain mouse monoclonal (WB: 1:1000; Santa Cruz Biotechnology sc-13524, RRID:AB_668849). Secondary antibodies used: HRP-conjugated goat anti-rabbit and mouse (WB: 1:50,000; Millipore 12-348 and 12-349, RRIDs:AB_390191 and AB_390192); Cy5-conjugated donkey anti-chicken, mouse, and goat (IF: 1:250; Jackson ImmunoResearch Labs 703-175-155, 715-175-150, and 705-175-147, RRIDs: AB_2340365, AB_2340819, and AB_2340415); DyLight 405-conjugated donkey anti-chicken (IF: 1:250; Jackson ImmunoResearch Labs 703-475-155, RRID: AB_2340373).

### DRG cultures

Cultures were generated from 4-6 month old Lis1 KO and no Cre control mice. Two methods were used to induce recombination. In some experiments mice were exposed to the 2 × 8 mg tamoxifen IP regimen then neurons harvested on day 4 after the first injection. In other experiments, 4-hydroxytamoxifen (4OHT, 2 µM) was added directly to the DRG cultures with these same genotypes without previous IP injections. Approximately 20 DRGs per mouse were harvested, then dissociated in Type XI collagenase (Sigma) for 1 hour at 37°C, followed by trituration with a flamed Pasteur pipet. Dissociated ganglia were then incubated in 0.05% trypsin (Invitrogen) at 37°C for 15 min. After a second trituration, the cell suspensions were centrifuged through a 12.5% BSA solution to remove myelin fragments and glial cells. Cells were then plated onto sterile, German glass coverslips (Fisher) coated with 10 μg/ml poly-D-lysine (>300 kDa; Sigma) and 10 μg/ml laminin (Millipore). Cells were cultured in DMEM/F12 medium (Corning) with 25mM HEPES, GlutaMAX (Thermo Fisher), N-2 supplement (Life Technologies), and 10% horse serum (HyClone). After 4 days in culture cells were either fixed for analysis of axon length and number of varicosities, or transferred to a live-cell imaging chamber for analysis of organelle transport.

### Analysis of axonal varicosities in DRG neurons

Neurons were cultured for 4 days, and then were fixed in 4% paraformaldehyde. Coverslips were mounted on glass slides using Prolong Gold Antifade (LifeTechnologies). Coverslips were imaged using the ImageXpress XLS (Molecular Devices) high content imaging system equipped with a 20X objective. Axon length and varicosity number were quantified using ImageJ software. The segmented line tool (1 pixel width) was used to trace the axon. Swellings that protruded visibly beyond the 1 pixel line on both sides were counted as varicosities.

### Analysis of organelle movement in DRG axons

Cultures grown on coated glass coverslips were exposed to 100 nM Lysotracker-Red (Millipore Inc.) for 20 min. Coverslips were transferred into fresh medium containing OxyFluor (Oxyrase Inc.) and 25 mM HEPES [pH 7.4], and placed in a custom-built water-heated microscope stage warmed to 37°C. Organelles were imaged using a Zeiss Axiovert 200 microscope equipped with a C-Apo 63x/1.2 W/0.2 water-immersion objective. Images were acquired at 0.5 sec intervals for 2 min using a Zeiss AxioCam HRm charge-coupled camera and linked AxioVision 4.7 software. Kymographs were generated from time-lapse movies using Image J software. Directionality of movement was determined by locating the relevant cell body prior to imaging. Lines showing a net displacement of ≥ 5 μm towards the cell body were categorized as retrograde, and their percentage (of all lines in the kymographs) were determined.

### Immunofluorescence

Tissue cryosections or cells were permeabilized in PBS with 0.1% Triton-X100 for 30 min, followed by incubation in blocking buffer consisting of 3% BSA (Fisher), 10% normal goat serum (Sigma), 0.2% Tween-20 (Biorad), in PBS. Samples were then incubated in primary antibody for 1 hour (cultures and nerve cross sections) to overnight (brain and spinal cord sections) at 4°C. After washing three times in PBS with 0.1% Tween-20, samples were incubated in fluorophore-conjugated secondary antibodies for the appropriate species for one hour at room temperature, washed, and mounted using Prolong Gold Antifade.

### Quantifying chromatolysis in brainstem sections

Coronal cryosections of mouse brainstem from animals exposed to the 2 × 8 mg tamoxifen regimen were stained with 1% toluidine blue. The Allen Mouse Brain Atlas was used as a guide to select coronal brainstem sections in which the nucleus ambiguus and other cardiorespiratory centers were likely located. Landmarks such as the fourth ventricle and pyramus granular layers were used to identify the proper sections. Comparisons were made from matched sections. Substantial Act-Cre-ER mediated recombination (GFP expression) was consistently observed in similar sections following tamoxifen administration. Two hallmarks of chromatolysis were examined – nuclear enlargement and nuclear acentricity. Image J was used to determine nuclear and somal areas and centroids. The ratio of the nuclear area to the somal area was calculated to establish a ‘nuclear enlargement index’. A ‘centroid displacement index’ was calculated by summing the X and Y displacement differences between the nuclear centroid and the somal centroid of each cell.

### RNA extraction and analysis

RNA was isolated from flash frozen tissues (see above) using the QIAGEN RNAEasy kit per the manufacturer’s instructions. RNA concentration was determined by fluorimetry using Ribogreen reagent (Thermo Fisher). 100 ng RNA was reverse transcribed with a SensiFast cDNA Synthesis Kit (Bioline). These cDNAs were used for quantitative droplet digital PCR (ddPCR) with Evagreen detection reagent and ddPCR Supermix (BioRad). Droplets for ddPCR were made using a QX200 Droplet Generator (Biorad). Results were analyzed using Poisson distribution on the QX200 Droplet Reader. Mouse Lis1 primer sequences were designed using NCBI BLAST (NCBI accession #NM_013625), and Harvard Primer Bank (https://pga.mgh.harvard.edu/primerbank/) was used for β-2 microglobulin primers (B2M; NCBI accession # NM_009735): Lis1 – forward, 5’ GCGAACTCTCAAGGGCCATA 3’ and reverse, 5’ CATTGTGATCGTGACCGTGC 3’; B2M – forward, 5’ TTCTGGTGCTTGTCTCACTGA 3’ and reverse, 5’ CAGTATGTTCGGCTTCCCATTC 3’.

### Experimental Design and Statistical Analysis

**Mice**: were treated with tamoxifen at 2-5 months of age and analysis of tissues was performed when mice showed symptoms of Lis1 KO. In all, 84 Lis1 KO animals were given the 2 × 8 mg tamoxifen regimen. 100% of these animals began to exhibit neurological symptoms (leg clasping, kyphosis, decreased motility) within a week. Of these, 18 died during that time, and 66 were euthanized when symptoms became severe. 38 no flLis1 control and 73 no Cre control mice were also given the 2 × 8 mg tamoxifen regimen. None of the controls showed any evidence of neurological disorder or malaise, but most were euthanized at the same time as the Lis1 KO animals to compare results. However 6 each of the control strains were monitored for 4 weeks after 2 × 8 mg tamoxifen and showed no symptoms. We also carried out mock injections of vehicle alone in 12 Lis1 KO animals. These controls also showed no symptoms.

### Culture Analyses

Axonal varicosities: DRGs cultures from 3 no Cre control and 3 Lis1 KO mice. 45 mm total axon length analyzed from control and 27 mm from Lis1 KO cultures. Transport: DRGs cultures from 2 no flLis1 control and 2 Lis1 KO mice; 27 axon segments (100 µm) analyzed per genotype. A total of 521 control organelles and 699 Lis1 KO organelles were included. Brainstem sections from 3 no flLis1 control and 4 Lis1 KO mice were used in the chromatolysis study. This includes analysis of 331 control neurons and 583 Lis1 KO neurons.

### Protein and RNA studies

Immunoblots are representative of typical results; little Lis1 protein reduction was observed in the cortex, while substantial, but sometimes variable, levels of reduction were observed in brainstem and cerebellum. The RNA analyses are from 3 mice for each treatment and genotype.

**Table 2:**
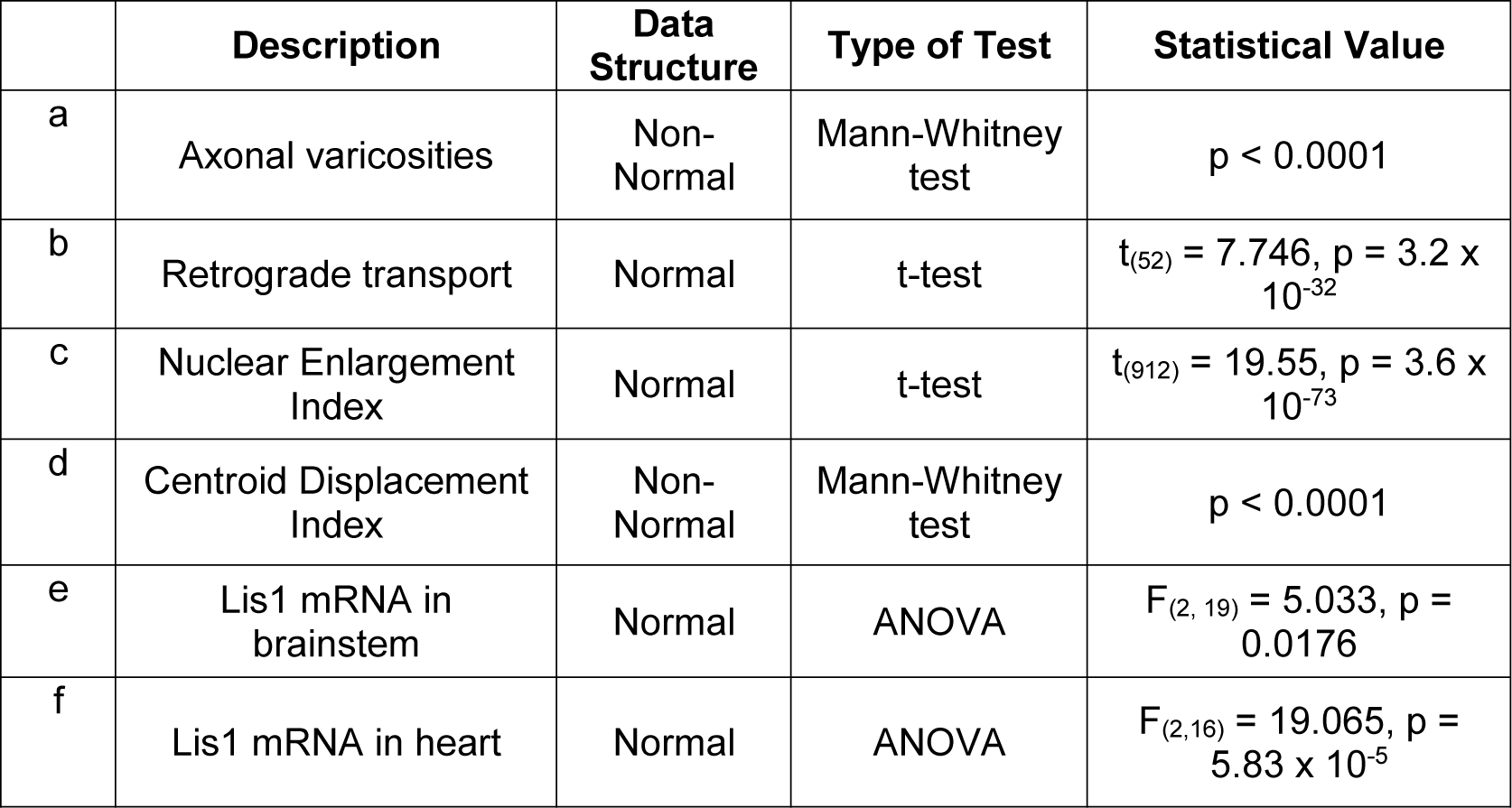
Statistical table.

|  | Description | Data Structure | Type of Test | Statistical Value |
| --- | --- | --- | --- | --- |
| a | Axonal varicosities | Non-Normal | Mann-Whitney test | $p < 0.0001$ |
| b | Retrograde transport | Normal | t-test | $t_{(52)} = 7.746, p = 3.2 \times 10^{-32}$ |
| c | Nuclear Enlargement Index | Normal | t-test | $t_{(912)} = 19.55, p = 3.6 \times 10^{-73}$ |
| d | Centroid Displacement Index | Non-Normal | Mann-Whitney test | $p < 0.0001$ |
| e | Lis1 mRNA in brainstem | Normal | ANOVA | $F_{(2, 19)} = 5.033, p = 0.0176$ |
| f | Lis1 mRNA in heart | Normal | ANOVA | $F_{(2, 16)} = 19.065, p = 5.83 \times 10^{-5}$ |

## RESULTS

### Lis1 is expressed in adult mouse tissues, prominently in the nervous system

Lis1 protein was found to be expressed in a number of adult mouse tissues. By automated capillary immunoblotting, levels of Lis1 protein were only modestly lower in adult brain than embryonic brain protein samples (Figure 1). Substantial Lis1 was also observed in adult spinal cord and dorsal root ganglia. While Lis1 could be detected in adult heart, liver, kidney, lung, and diaphragm, the levels were much lower in all non-nervous tissues tested than in brain, spinal cord and DRG. Therefore, Lis1 may function in other tissues, but is either present in fewer cells or at lower amounts per cell.

### Knockout (KO) of Lis1 by Cre-mediated recombination causes axon transport defects in cultured adult sensory neurons

All mice strains used in these studies are described in table 1 in the methods section. To test the effectiveness of the Act-Cre-ER model for inducing recombination in neurons and reducing Lis1 expression, DRGs from Lis1 KO mice and the no Cre controls were dissociated and cells cultured for 24 hours. Cultures were treated with 4-OHT for an additional 72 hours. No tamoxifen-induced recombination was observed in the no Cre control cultures, as detected by prominent tdTomato fluorescence but no GFP (Figure 2A). In contrast, substantial recombination was observed in the majority of neurons in the Lis1 KO cultures, detected by the presence of bright GFP fluorescence (Figure 2B). GFP was observed in both axons and cell bodies. Also, Lis1 protein levels in extracts prepared from Lis1 KO cultures was greatly reduced relative to no Cre control cultures (Figure 2C). Together these findings demonstrate effective KO of Lis1 by bath-applied 4-OHT in cultures of Lis1 KO mice but not no Cre controls.

IP injection of tamoxifen into adult Lis1 KO mice also resulted in recombination in DRGs. For this experiment we injected 8 mg of tamoxifen on two consecutive days (2 × 8 mg regimen – see methods) and harvested DRGs on day 4 after the first injection. Bright GFP fluorescence was observed in neuronal plasma membranes and satellite cells in DRG sections from these animals (Figure 2D). No GFP was detected in DRGs from no Cre control animals injected at the same time (not shown). Dissociated cultures prepared from no Cre control DRGs showed only tdTomato fluorescence (Figure 2E), while Lis1 KO cultures exhibited bright GFP fluorescence indicative of substantial recombination (Figure 2F). We immunostained these cultures for neurofilament (NF) to label axonal processes specifically so that images could be analyzed using automated software algorithms. However, NF staining was different in the no Cre control and Lis1 KO neurons (Figure 2E, F). NF was uniformly distributed along axonal processes in no Cre control neurons (Figure 2E), while in Lis1 KO neurons NF was more prominent in axon endings, possibly reflecting altered neurofilament transport (Figure 2F). Visual inspection and manual measurements in ImageJ showed Lis1 KO neurons had fewer and shorter axons (not shown) and also had significantly more varicosities than the no Cre controls (Mann-Whitney test, p < 0.001^a^; Figure 2G). Although varicosities occur normally at sites of growth cone pausing or sites of branch formation (Rees et al., 1976; Malkinson and Spira, 2010) an increased number is often associated with axonal blockages due to transport defects (Liu et al., 2012). Indeed, reduced retrograde transport of acidic organelles in living axons was observed in Lis1 KO axons compared to no Cre controls (t-test, p = 3.2 × 10^−32^ ^b^; Figure 2H) consistent with other studies in adult rat DRG neurons where Lis1 was depleted using siRNA transfections (Pandey and Smith, 2011).

### Lis1 KO causes a severe phenotype in adult mice

Surprisingly, the 2 × 8 mg regimen caused a rapid decline in health of Lis1 KO animals, with spinal kyphosis (Figure 3A) and hind leg clasping (Figure 3B) observed 4 days after the first injection. None of the control animals showed this phenotype. In early experiments animals died within a week after the first injection, and mice were subsequently euthanized as soon as they began to exhibit symptoms, typically on days 3-5. Animals that were given a different regimen of tamoxifen, 2 mg injected for 5 consecutive days (5 × 2 mg regimen), remained non-symptomatic for nearly two weeks, after which they exhibited similar symptoms as observed in the 2 × 8 mg regimen. Figure 3C shows symptom-free survival duration plots for these animals. No differences were observed between male and female animals. Most animals were 2 months old at the time of injection, but similar responses were observed in older animals (4-5 months). Control animals did not exhibit any symptoms but were typically euthanized at the same time as KOs to be able to compare tissues for extent of recombination and Lis1 expression levels. However, 6 no Cre control and 6 *CAG-cre/Esr1* animals that received the 2 × 8 mg tamoxifen regimen lived for over a month with no detectable symptoms (Figure 3C).

**Figure 3.**
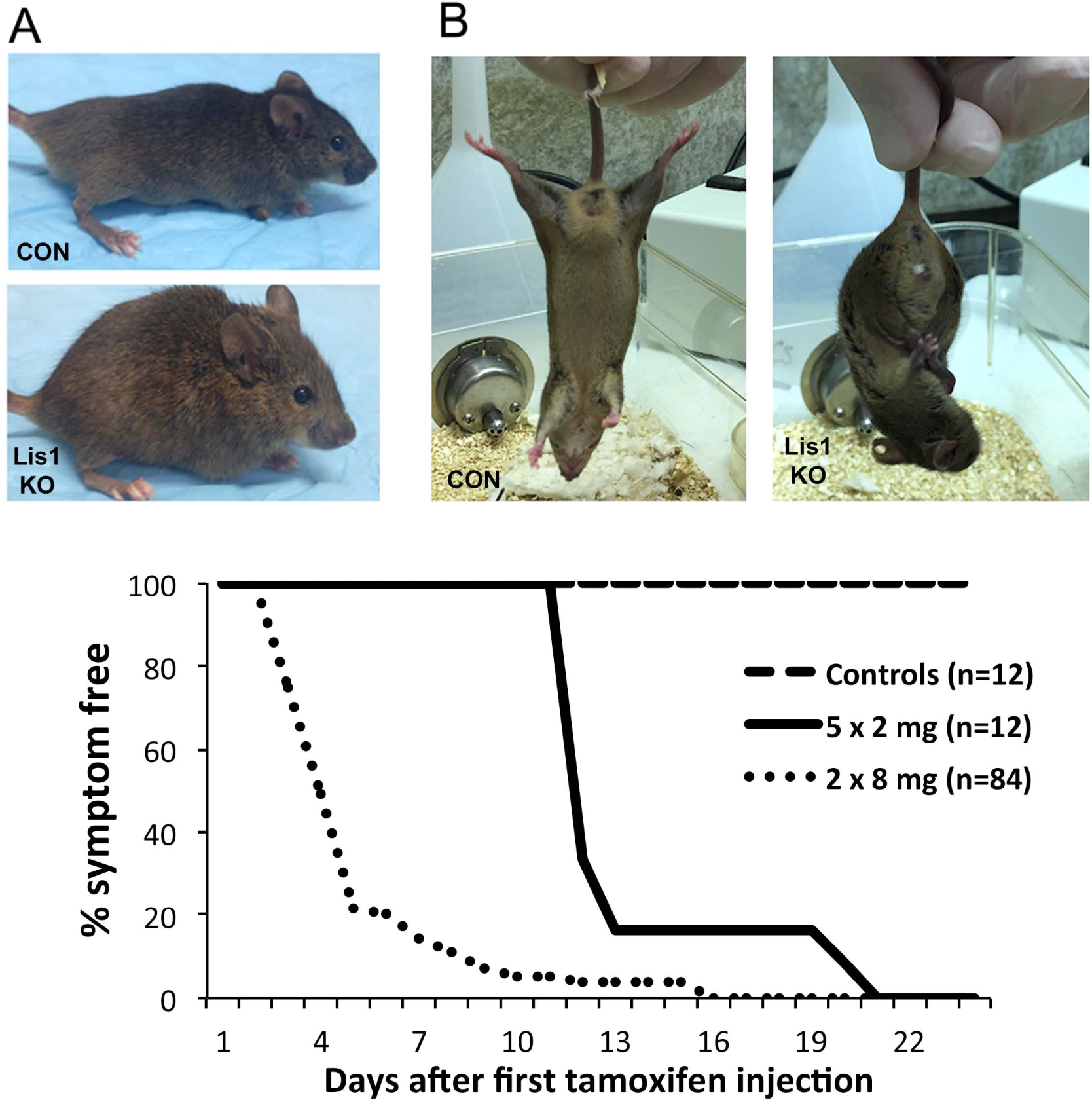
Lis1 KO via IP tamoxifen injection in adult mice results in severe malaise. Lis1 KO mice exposed to tamoxifen invariably displayed spinal kyphosis **(A, lower panel)** and hind leg clasping **(B, right panel)**. Neither symptom was observed in control animals (CON) at any time. Controls in A and B are no Cre control, but no symptoms were observed in three other control groups no flLis1, Lis1 KO het, or mock-injected Lis1 KO animals. Mice were euthanized as soon as kyphosis and leg clasping became apparent. These developed with different latencies in Lis1 KO mice depending on the specific tamoxifen-dosing regimen. **C) S**ymptom-free survival curves show that the latency is shorter for the 2 × 8 mg regimen compared to the 5 × 2 mg regimen (see methods). Control mice were typically euthanized at the same time as the Lis1 KO mice for recombination and expression studies. However, 6 no Cre control mice and 6 no flLis1 control mice receiving the 2 × 8 mg dosing regimen survived symptom free for 3 weeks before they were euthanized.

### Temporal differences in tamoxifen-induced recombination in distinct brain regions of Lis1 KO mice

Lis1 KO mice and no Cre control strains were given the 2 × 8 mg tamoxifen regimen. As expected, the no Cre control brains only exhibited tdTomato fluorescence on day 4 after the first injection (Figure 4A, left). In contrast, bright GFP fluorescence was detected in Lis1 KO mice, but surprisingly, primarily in midbrain and hindbrain regions, with little fluorescence detected in the cortices (Figure 4A, right). A similar result was observed in no flLis1 control so the recombination pattern is similar regardless of whether or not the mice carry the floxed Lis1 alleles.

**Figure 4.**
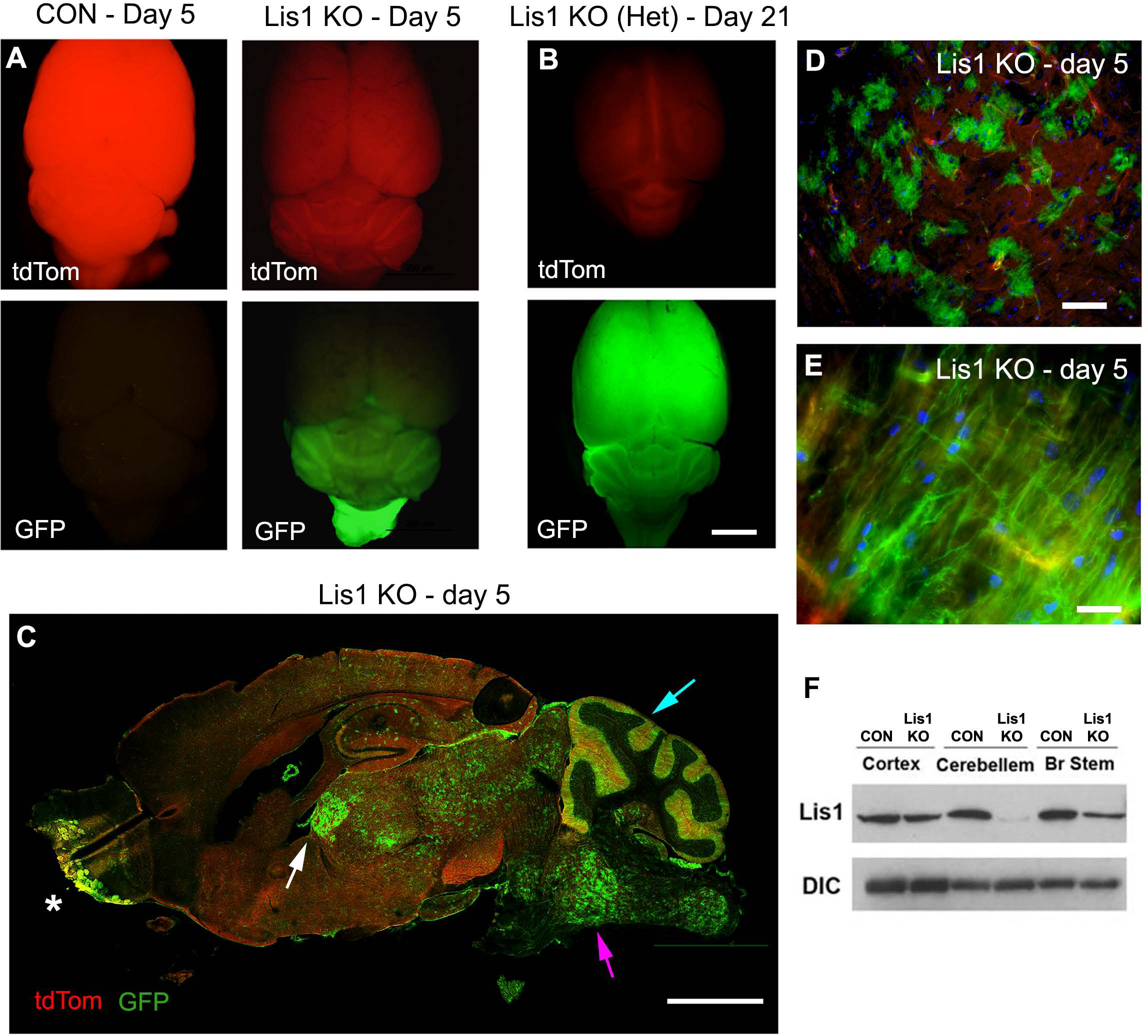
Cre-dependent recombination in the brain after tamoxifen injection. **A)** On day 5 after the 2 × 8 mg tamoxifen regimen a no Cre control mouse (CON - day 5) had bright red fluorescence (top left panel) but showed no GFP fluorescence indicative of recombination (lower left panel). This animal showed no sign of neurological problems. A Lis1 KO mouse (Lis1 KO - day 5) showed reduced red fluorescence (top right panel) and GFP primarily in the hindbrain (lower right panel). This animal exhibited kyphosis and leg clasping. **B)** Lis1 KO het mouse (Lis1 KO (Het) - day 21) showed no sign of neurological problems, and was euthanized for tissue collection on day 21. GFP expression was observed throughout the brain. **C)** A sagittal section through the brain of a Lis1 KO mouse on day 5 (Lis1 KO - day 5) shows mosaic recombination enriched in midbrain (white arrow), hindbrain (magenta arrow), and cerebellum (blue arrow), with widely scattered cells in cortex and hippocampus. Recombination also occurs in olfactory bulb (asterisk). **D)** Using higher magnification GFP-positive cells in the midbrain can be seen interspersed with cells that have not yet undergone recombination. **E)** Fibers labeled with GFP are clearly visible in the brainstem. **F)** Lis1 expression is reduced in extracts from brainstem and cerebellum of Lis1 KO mice compared to no Cre controls. Scale bars: A, B and C = 5 mm; D= 100 µm; E = 20 µm.

Although the severity of the phenotype in Lis1 KO mice prevented examination of recombination at later times, no flLis1 control animals Lis1 KO het that remained symptomless for at least 3 weeks after the 2 × 8 mg tamoxifen regimen. These animals showed substantial recombination observed across the entire brain, including the cortex, demonstrating variable rates of recombination in different brain regions with this 2 × 8 mg tamoxifen regimen (shown for Lis1 KO het in Figure 4B).

In Lis1 KO mice recombination was observed throughout the medulla pons, midbrain and cerebellum and into the spinal cord, but only sparsely in the cortex and hippocampus (Figure 4C). At this level of analysis, the most prominent recombination in the cerebellum occurs in the molecular layer in linear profiles reminiscent of Bergmann's glia. Substantial recombination was also observed in the olfactory bulb and choroid plexus (Figure 4C). GFP positive cells in the brainstem appeared stellate in shape (Figure 4D). White matter tracks in the brainstem and cervical spinal cord were also GFP positive (Figure 4E). This GFP distribution correlates with the reduced Lis1 expression observed in brainstem and cerebellum but not in cortex (Figure 4F).

### Lis1 KO occurs in both neurons and glia

In cryosections of Lis1 KO DRGs (2 × 8 mg regimen), recombination was observed in both neurons and satellite cells (Figure 2C). Membrane-targeted GFP was observed along neuronal membranes in the brainstem, Purkinje cells in the cerebellum, and motor neurons in the spinal cord (Figure 5 A-F). It was more difficult to distinguish neuronal processes from glia or axons from dendrites in the neuropil. Since Lis1 depleted DRGs neurons show altered axon transport and DRGs cultured from Lis1 KO animals showed signs of altered axon transport, we examined cross sections of phrenic, vagus, and sciatic nerves, as well as spinal cord ventral roots. Two concentric rings of GFP were typically observed around myelinated axons (Figure 5G-J). The outer ring flanked the periphery of the myelin sheaths (Figure 5I) that stained for myelin basic protein (Figure 5J). GFP was not observed in the tightly packed myelin sheath itself, so the outer ring likely represents the plasma membrane of the myelinating glial cell. The inner ring was juxtaposed along the axoplasmic membrane of the ensheathed axon, and we interpret this as representing tamoxifen-induced recombination in axons of Lis1 KO neurons. This interpretation is strengthened by the observation that only an outer GFP ring was observed in some myelinated axons (Figure 5I), which would be unlikely if the inner GFP-positive ring was also part of the myelinating Schwann cell. Unmyelinated C-fiber bundles in the sciatic nerve are ensheathed by membranes of Schwann cells that do not form myelin. These “Remak bundles” can be observed by EM (Figure 5K). Red and green fluorescent rings were often observed in the same Remak bundles indicating that some, but not all axons in the Remak bundle had undergone recombination (Figure 5L). Together these data support the *in vitro* finding that recombination, and by inference, Lis1 KO, occurs in both neurons and glia. The preponderance of recombination in the midbrain, hindbrain, and PNS, coupled with reduction in Lis1 protein levels in these regions, suggest that Lis1 KO in neurons and glia in these regions contributes to the observed neurological phenotypes.

**Figure 5.**
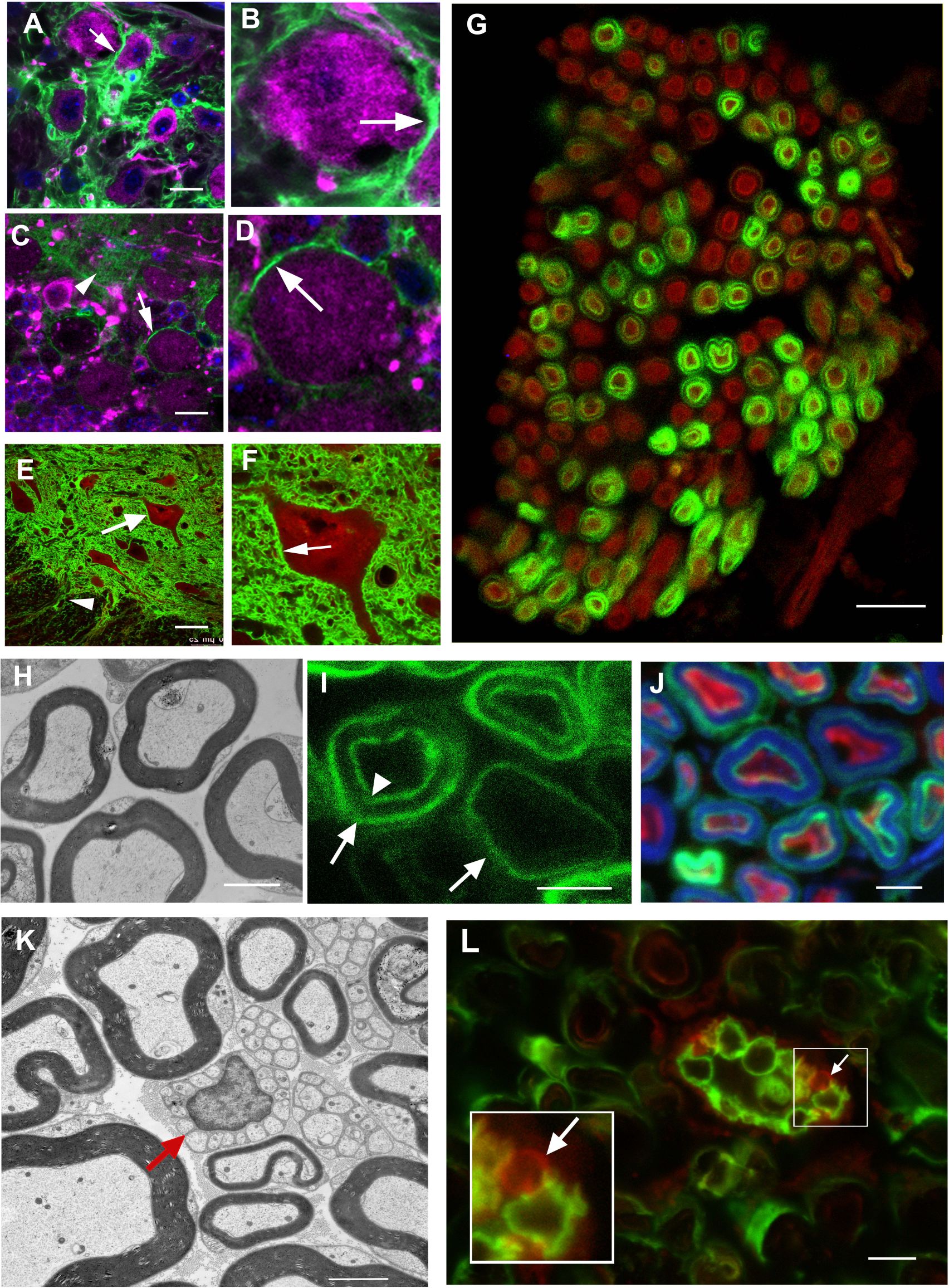
Both neurons and glia show evidence Cre-dependent recombination. **A)** GFP-positive neuropil surrounds MAP2-positive neurons (magenta) in a brainstem region thought to be the nucleus ambiguus in a mouse exposed to the 2 × 8 mg tamoxifen regimen. **B)** The neuron indicated in A has been digitally enlarged to show details. The arrow points to possible neuronal plasma membrane. **C)** Purkinje cells in the cerebellum stained with MAP2 (magenta) also have GFP-positive plasma membranes (white arrow) indicating that recombination occurred in these neurons. Neuropil in the molecular layer is also GFP positive (arrowhead). **D)** The neuron in C has been digitally enlarged to show detail. **E)** GFP-positive neuropil surrounds a motor neuron (white arrow) labeled with ChAT (red) in the anterior horn of the thoracic spinal cord. GFP positive fibers (arrowhead) can be seen coursing towards the ventral root. **F)** The motor neuron indicated in E is digitally enlarged to show detail. The white arrow points to apparent neuronal plasma membrane. **G)** A cross section through the phrenic nerve shows concentric rings of GFP around approximately half of the neurofilament-positive axons (red). **H)** EM of a cross section through a WT mouse nerve showing myelinated axons. **I)** GFP can be observed as two concentric rings or single rings (arrows, outer ring, arrowhead, inner ring). **J)** The area between concentric rings is positive for myelin basic protein (blue), while the inside of the inner ring is positive for NF (red). **K)** EM of a cross section of WT mouse sciatic nerve showing Remak bundles of unmyelinated axons surrounded by a single glial cell (red arrow). **L)** Cross section of sciatic nerve from Lis1 KO mouse with a Remak bundle containing some GFP-encircled axons, and some without encircling GFP (arrow – positive for tdTomato only). Inset is digitally enlarged to show an axon without recombination (red, arrow) alongside recombined axons (green). Scale bars: A, C, G = 10 µm; E = 30 µm; H-L = 2 µm.

### Brainstem neurons show chromatolysis in Lis1 KO mice

Figure 6A shows GFP expression in a coronal section through the hindbrain, with a dense concentration of GFP-positive cells in the ventral hindbrain. This area contains nuclei that are vital for cardiorespiratory function, and complete functional loss of these neurons would result in rapid death (Melov et al., 1998; Quintana et al., 2012). Impairment of axonal transport or other dynein-dependent processes in this region could account for the severe phenotype in Lis1 KO animals. Experimental axotomy and diseases that involve axonal dysfunction can produce the cell body response of chromatolysis (Cragg, 1970; Hanz and Fainzilber, 2006). Though the mechanisms underlying chromatolysis remain hypothetical, the process is characterized by nuclear swelling and nuclear acentricity, both of which were observed in GFP-rich regions in coronal sections through the brainstem of Lis1 KO (Figure 6C). The nuclei were significantly larger and more acentric than controls providing evidence for axonal dysfunction in these neurons (t-test, p = 3.6 × 10^−73^ ^c^ (D); Mann-Whitney test, p < 0.0001 ^d^ (F); Figure 6D-G).

**Figure 6.**
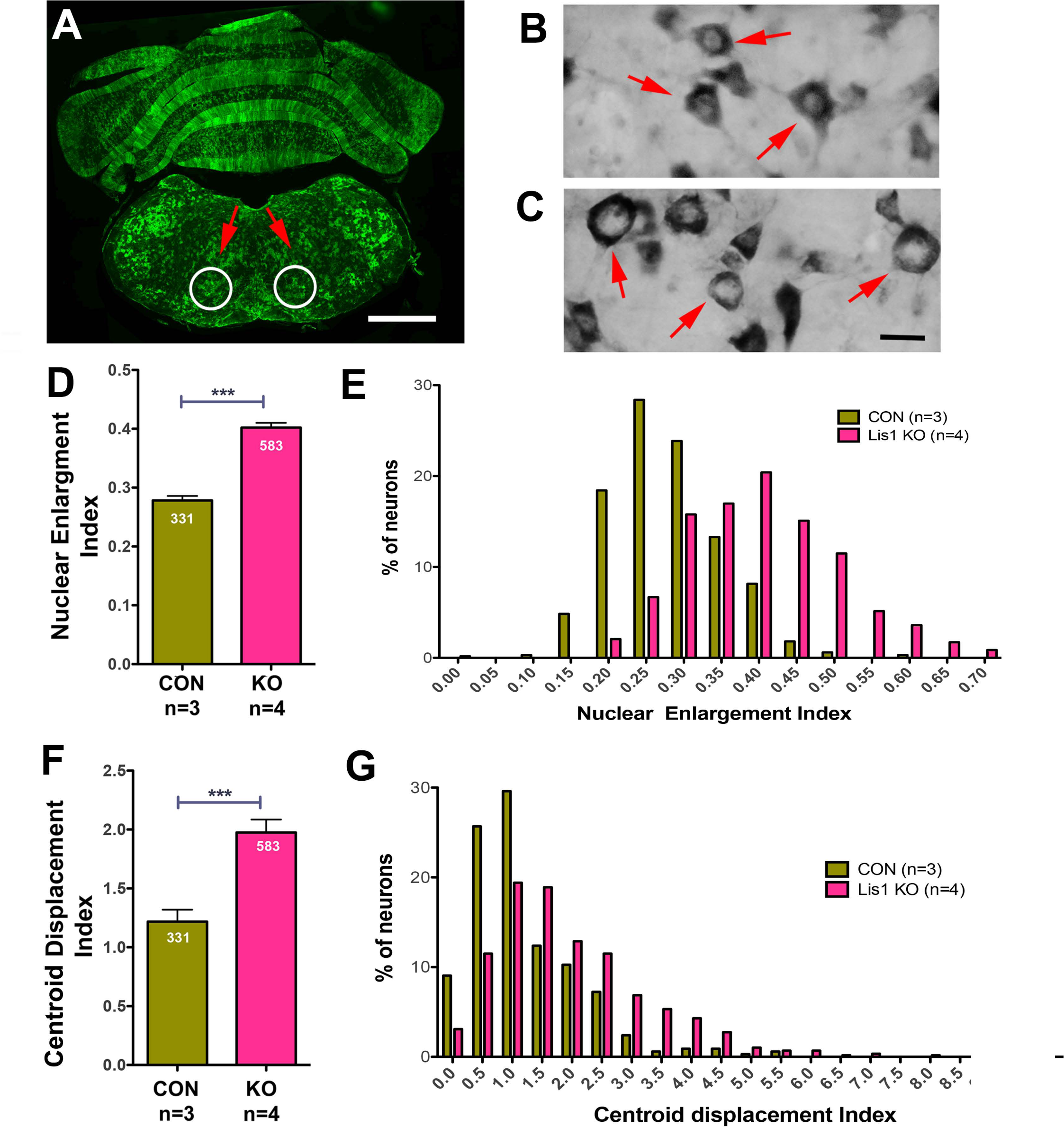
Brainstem neurons in Lis1 KO mice exhibit signs of chromatolysis. **A)** A coronal section through the hindbrain on day 4 after the 2 × 8 mg tamoxifen regimen shows extensive recombination in the ventral brainstem containing cardiorespiratory centers. White circles indicate the region used in the analyses of chromatolysis. **B, C)** Sections were stained with toluidine blue to determine the size and position of the nucleus in neurons in the indicated regions. The neurons in B are from a no flLis1 control mouse. The neurons in C are from a Lis1 KO animal. **D)** A nuclear enlargement index, (see methods) was used to compare nuclear enlargement in no flLis1 controls (CON) and Lis1 KO (KO). **E)** The histogram shows the distribution of this index in CON and Lis1 KO neurons. **F)** The position of the nucleus within the soma was also determined using the centroid displacement index (see methods). This involves determining the centroid position of both the nucleus and soma and calculating the total displacement distance (µm) of the nuclear centroid from the somal centroid. **G)** Histogram showing the distribution of CDI found in CON and KO neurons. The number of neurons measured and the number of mice (n) are indicated for **D-G.** Bars indicate mean ± 95% CI. Significance determined by students t-test (D), Mann-Whitney test (F). *** p<0.001. For details see Results. Scale bars: A = 1 mm; B, C = 10 µm.

### Lis1 loss in the hindbrain is the most likely cause of the KO phenotype

Lis1 KO mice receiving the 2 × 8 mg tamoxifen regimen had a much more rapid onset of symptoms compared to the 5 × 2 mg regimen (Figure 3C). Indeed, at a time when the 2 × 8 mg mice were severely affected (day 5), the 5 × 2 mg animals had no overt symptoms. Substantially more recombination was observed in the brains of 2 × 8 mg animals compared to 5 × 2 mg animals, which correlates with the onset of severe symptoms (Figure 7A) and the level of Lis1 mRNA in the brainstem (one-way ANOVA, F_(2,19)_ = 5.033, p = 0.0176^g^; Figure 7B). In contrast, recombination in the heart was similar in both sets of mice (Figure 7C), as were Lis1 mRNA levels, which were reduced equally with both regimens (one-way ANOVA, F_(2,16)_ = 19.065, p = x 10^−5^ ^h^; Figure 7D). This indicates that Lis1 KO in the heart is less likely to be responsible for the early onset of symptoms. Other tissues with sporadic GFP (lung, liver, and kidney) also showed a similar degree of recombination with both regimens, supporting the idea that the hindbrain loss of Lis1 contributes significantly to the KO phenotype.

**Figure 7.**
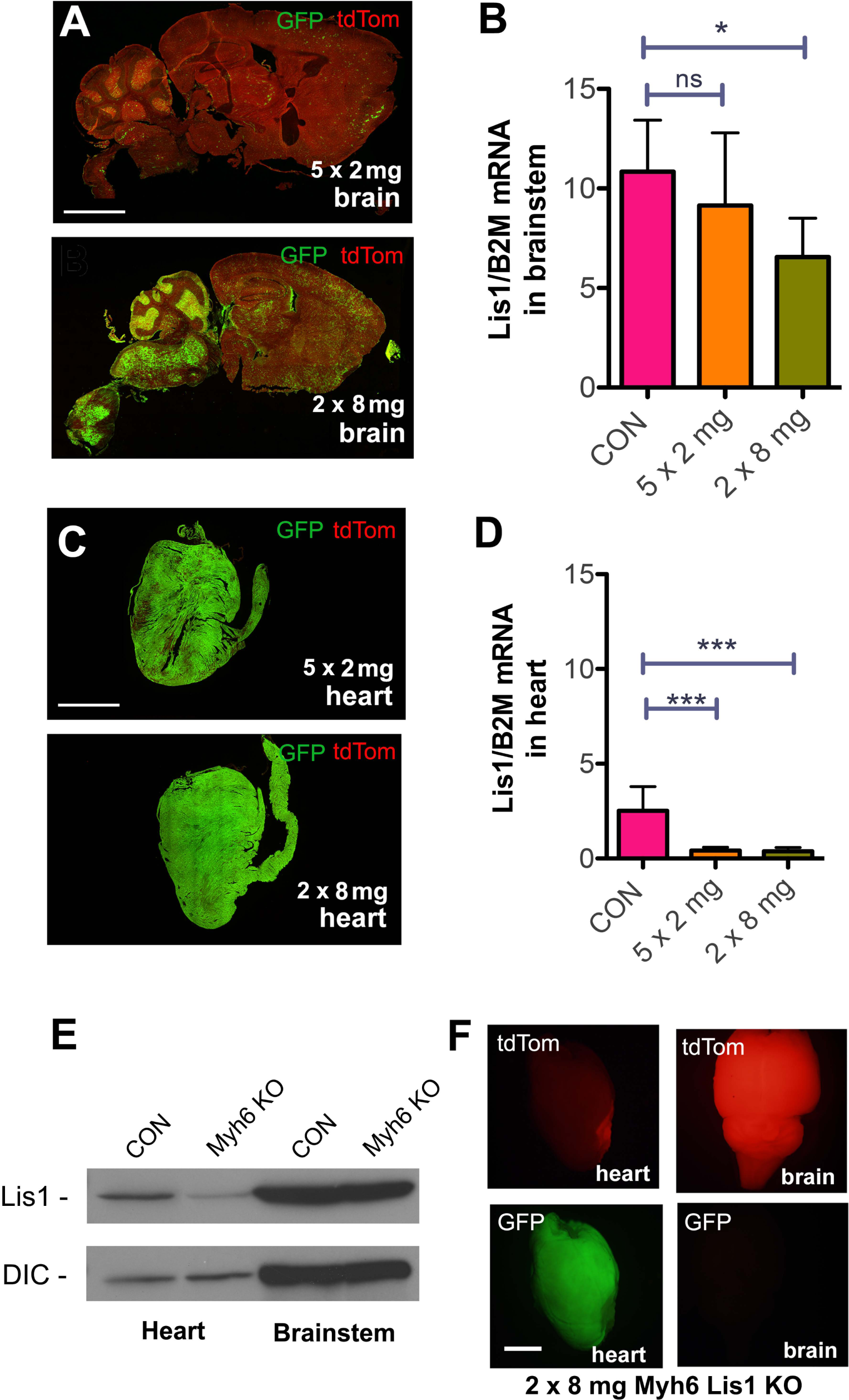
Comparing the effect of Lis1 KO in brainstem and heart. **A)** Sagittal brain sections of Lis1 KO mice 5 days after the initial injection of either 5 injections of 2 mg tamoxifen (top) or 2 injections of 8 mg tamoxifen (bottom). The 2 × 8 mg treatment resulted in much higher GFP expression than the 5 × 2 mg treatment, particularly in the brainstem and cerebellum. **B)** Lis1 mRNA levels normalized to β2 microglobulin (B2M) mRNA levels from brainstem of no Cre control mice injected with 2 × 8 mg tamoxifen (CON), and Lis1 KO mice injected with either 5 × 2 mg or 2 × 8 mg tamoxifen. Lis1 mRNA levels were significantly decreased in brainstem of 2 × 8 mg animals, but not 5 × 2 mg animals, relative to no Cre controls, 5 days after initial injection. **C)** Sections of heart from 5 × 2 mg (top) and 2 × 8 mg (bottom) treated Lis1 KO mice. Both the 2 × 8 mg and 5 × 2 mg treatment resulted in similar levels of GFP expression in heart. **D)** Lis1 mRNA levels normalized to B2M mRNA levels from heart of 2 × 8 mg injected no Cre control (CON), 5 × 2 mg, and 2 × 8 mg treated mice. Lis1 mRNA levels were reduced significantly in both the 5 × 2 mg and 2 × 8 mg treated mice relative to the no Cre control, but were not significantly different from each other. **E)** Western blot of brainstem and heart lysates from cardiomyocyte-specific Myh6 KO mice show reduced levels of Lis1 protein in heart, but not brainstem compared to no Cre control mice (CON). Dynein intermediate chain (DIC) was used as a loading control. **F)** Whole mount brain (right) and heart (left) from Myh6 KO mouse show recombination (GFP) in heart, but not brain. Significance in B and D determined by one way ANOVA. * p<0.05, *** p< 0.001. For details see results. Scale bars in A and C – 5 mm; F – 2 mm.

We performed another experiment to directly test contributions of Lis1 KO in the heart to the phenotype observed in Lis1 KO mice by generating an inducible KO in which Cre-ER is driven by a Myh6 promotor (Myh6 KO) (Sohal et al., 2001). As expected, tamoxifen injections (2 × 8 mg) reduced Lis1 expression in the heart but not in the brainstem of these mice (Figure 7E). Moreover, robust GFP expression, and thus Myh6 KO-dependent recombination, was observed in the heart, but not in the brain (Figure 7F). Despite significant recombination in the heart, 0/12 mice showed any detectable phenotype, and all lived until apparently symptom free, until euthanized 4 weeks later. Together these data provide evidence that loss of Lis1 in midbrain/hindbrain neurons is responsible for severe phenotype in Lis1 KO mice.

## Discussion

In this study we show that Lis1 KO by tamoxifen-induced recombination in Lis1 KO mice causes neuropathology and lethality. Cultured DRG neurons from Lis1 KO animals have axon transport defects and axonal varicosities. Given the preponderance of evidence that Lis1 regulates cytoplasmic dynein, along with the temporal and spatial pattern of recombination and evidence of chromatolysis, the simplest explanation for the severe phenotype is that defective axon transport causes pathological changes in critical subsets of neurons in the brainstem. Lis1 protein is expressed at higher levels in adult brain than in other tissues, and KO specifically in the heart produced no obvious phenotype. However, Lis1 may also play important but undiscovered roles in other tissues.

The finding that Lis1 continues to play a vital role after the majority of mitosis and migration in the brain has occurred supports the idea that Lis1 regulates other dynein-dependent processes like axonal transport. Mouse models in which Lis1 is knocked out developmentally have revealed a much more severe phenotype in homozygotes (preimplantation lethality) relative to heterozygotes (developmental delay in some regions) (Hirotsune et al., 1998). Our data indicate that this dosage effect is retained in mature animals. Evidence of chromatolysis in brainstem neurons, along with defective morphology and organelle motility in DRG neurons from Lis1 KO mice adds credence to a proposed role for Lis1 in axonal transport and vesicle dynamics in adult mammals.

While our studies unequivocally demonstrate a vital role for Lis1 in adult mice, there are still many unanswered questions. Dynein-based transport also occurs in dendrites, although this has not been studied to the same extent as dynein-based transport in axons (Tear, 2008; Zheng et al., 2008; Kapitein et al., 2010; Wiggins et al., 2012; Arthur et al., 2015; Ayloo et al., 2017). Lis1 reportedly impacts retrograde translocation of excitatory synapses in dendrites in hippocampal cultures and organotypic slice cultures from newborn mice (Kawabata et al., 2012) and Lis1 heterozygosity can influence spine morphology in adolescent mice and impacts social behavior (Sudarov et al., 2013), so it is very likely that postsynaptic processes are also altered in our Lis1 KO mice.

Lis1 is present in Schwann cells in adult DRG cultures, as well as in glial cells in cortical and hippocampal cultures (personal observation). Several reports indicate that Schwann cells and oligodendrocytes also utilize dynein during myelination (Gould et al., 2000; Langworthy and Appel, 2012; Yang et al., 2015). It is clear from our Lis1 KO reporter studies that glial cells exhibit recombination. If Lis1 contributes to dynein functions in these cells, perturbations in the mouse could affect myelin integrity and dynein-based transport in glial cells, indirectly disrupting neural circuitry. Generating glial and neuronal specific KO animals may allow us to tease apart the contribution of Lis1 to dynein function in different neural cell types.

While tamoxifen induced Lis1 KO in Lis1 KO mice generated a severe and readily observable phenotype, the choice of this Cre-driver complicated the neural specific analysis because other tissues were also targeted for recombination. The severe Lis1 KO phenotype was clearly not due to loss of Lis1 in cardiac muscle, but we cannot completely exclude contributions of Lis1 loss in other tissues. Notably, the recombination in these tissues was highly mosaic at the time when neurological symptoms were severe and euthanasia was performed (not shown). Urine and blood was tested for changes that might signal kidney, bladder, liver or intestine defects, but no significant changes were found (not shown). To try to limit Lis1 loss to the brain we performed stereotaxic injections of 4OHT into lateral ventricles in the Lis1 KO but this resulted in many fewer GFP-positive cells and did not produce a detectable phenotype (not shown). Thus, more selective examination of the role of Lis1 in adult neuronal circuits will require using Cre-driver(s) specific for different neuronal populations.

An unexpected outcome of our study was the temporal variability in onset of tamoxifen-induced recombination in different brain regions. No Cre control or Lis1 KO het mice remained healthy after the 2 × 8 mg tamoxifen regimen and had very widespread recombination throughout the brain three weeks after the first injection. However, by the time Lis1 KO mice exhibited severe symptoms, recombination was only prominent in the midbrain hindbrain and PNS, so Lis1 loss in the cortex and hippocampus was unlikely to contribute to the KO phenotype. Recombination was observed in axons of the vagus, phrenic and sciatic nerve as well as ventral roots at onset of severe neurological symptoms, adding support to the hypothesis that loss of Lis1 in cardiorespiratory neurons could be a major factor in the Lis1 KO phenotype. The vagus nerve includes autonomic axons emerging or converging on the nucleus ambiguus, solitary nucleus, dorsal nucleus of vagus, and spinal trigeminal nucleus and is involved in cardiorespiratory control. The phrenic nerve contains sympathetic, sensory and motor axons innervating the diaphragm, mediastinal pleura, and pericardium. Sensory and motor neurons with axons in sciatic nerve are less likely to contribute to the death of the animals, but may contribute to leg clasping and kyphosis observed in 100% of Lis1 KO mice.

Brainstem perturbations in animals can lead to death due to cardio-respiratory disruption (Talman and Lin, 2013; Piroli et al., 2016; Sun et al., 2017). Our data strongly suggests that Lis1 KO in brainstem neurons is a lethal event in mice. Our study raises the question of whether axonal transport is compromised in children with LIS, and if this could contribute to the increasingly frequent and severe seizures and early lethality exhibited by these patients. The brain malformations that occur in utero will not be easy to address in LIS patients. However, axon transport defects might be amenable to treatments after birth with drugs that target the dynein regulatory machinery. Moreover, the symptoms of Perry Syndrome suggest that there is at least respiratory involvement in the disorder(Wider and Wszolek, 2008). It remains to be determined if less dramatic alterations in the LIS1 gene that might cause more subtle defects in LIS1 expression or function contribute to other types of neurological disorders. There is a growing body of evidence that dynein-related proteins are linked to many neurodegenerative diseases, including SMA, Charcot-Marie-Tooth and Perry Syndrome (Rees et al., 1976; Wider and Wszolek, 2008; Weedon et al., 2011; Harms et al., 2012; Neveling et al., 2013; Oates et al., 2013; Peeters et al., 2013). Resolving the role of Lis1 in post-developmental neurons and elucidating the pathways regulating axonal transport could uncover new targets for treatment of these neurodegenerative conditions.

In order to treat disorders involving axons transport defects it is critical to elucidate all of the temporal and spatial regulatory mechanisms controlling the transport motors in axons. Many cell culture experiments indicate that Lis1 stimulates dynein processivity(Liu et al., 2000; Smith et al., 2000; Pandey and Smith, 2011; Shao et al., 2013; Klinman and Holzbaur, 2015; Villarin et al., 2016). However, one transport study indicated that Lis1 knockdown increased mitochondrial transport(Vagnoni et al., 2016), and several *in vitro* biophysical studies showed that Lis1 inhibited processivity of purified dynein (Yamada et al., 2008; McKenney et al., 2010; Huang et al., 2012). More recent assays using purified proteins are beginning to reveal how this might occur at the molecular level in the context of other dynein regulators like dynactin and BICD2 (Baumbach et al., 2017; DeSantis et al., 2017; Gutierrez et al., 2017). In those studies Lis1 dramatically increased dynein processivity. Interestingly BICD2 mutations that cause SMALED stimulate dynein processivity, so motor activity must be finely tuned (Huynh and Vale, 2017). Kinase pathways that impact dynein function have been previously identified. CDK5, mutations in which have been linked to LIS (Magen et al., 2015; Parrini et al., 2016) and CDK1 phosphorylate and regulate the Lis1- and dynein-interacting protein Ndel1 (Hebbar et al., 2008; Pandey and Smith, 2011). Insulin dependent inhibition of GSK3β, a kinase with a growing list of potential neurological disease links (Dell'Osso et al., 2016), phosphorylates dynein and regulates its interactions with Ndel1 and APC (Gao et al., 2015; Gao et al., 2017). It will be interesting to determine if these pathways can be manipulated to alter the severity of the Lis1 KO phenotype, and if they can be used in trying to alleviate symptoms of patients with diseases caused by transport defects.

## Acknowledgements

We would also like to acknowledge Tia Davis for her help with mouse colony maintenance.

